# Increased molecular plasticity along the septotemporal axis of the hippocampus is associated with network functioning

**DOI:** 10.1101/593095

**Authors:** Evangelos Sotiriou, Fevronia Angelatou, Costas Papatheodoropoulos

## Abstract

Molecular plasticity crucially supports adaptive cellular and network functioning in the brain. Variations in molecular plasticity may yield important differences in neuronal network dynamics between discrete brain subregions. In the present study we show that the gradual development of sharp waves (SPWs), a spontaneous network activity that is organized under normal *in vitro* conditions in the CA1 field of ventral but not dorsal hippocampal slices, is associated with region selective molecular reorganization. In particular, increased levels of mRNAs for specific GABA_A_ receptor subunits (α1, β2, γ2) occurred in ventral hippocampal CA1 field during the development of SPWs. These mRNA changes were followed by a clear increase in GABA_A_ receptor number in ventral hippocampus, as shown by [^3^H]muscimol binding. An increase in mRNAs was also observed in dorsal slices for α2 and α5 subunits, not followed by quantitative GABA_A_ receptor changes. Furthermore, full development of SPWs in the CA1 field (at 3 hours of slice maintenance *in vitro*) was followed by increased expression of immediate early genes c-fos and zif-268 in ventral hippocampal slices (measured at 5 hours *in vitro*). No change in c-fos and zif-268 levels is observed in the CA1 field of dorsal slices, which do not develop spontaneous activity. These results suggest that generation of SPWs could trigger specific molecular reorganization in the VH that may be related to the functional roles of SPWs. Correspondingly, the revealed increased potentiality of the ventral hippocampus for molecular reorganization may provide a clue to mechanisms that underlie the regulated emergence of SPWs along the longitudinal axis of the hippocampus. Furthermore, the present evidence suggests that dynamic tuning between spontaneous neuronal activity and molecular organization may importantly contribute to the functional segregation/heterogeneity seen along the hippocampus.

## Introduction

Molecular plasticity represents a fundamental parameter in brain functioning that permits rapid adaptational responses of neural circuits to current need of the organism, thereby supporting complex brain functions like learning and memory by providing the building blocks for the formation of lasting changes at higher levels of neural organization (Ziv and Brenner 2018; Minatohara et al. 2016; Diekelmann and Born 2010). Hence, mechanisms of molecular plasticity are essential in supporting maintained changes in synaptic transmission that underlie learning and memory (Shonesy et al. 2014; Blackwell and Jedrzejewska-Szmek 2013; Elgersma and Silva 1999; Colbran 2015; Kotaleski and Blackwell 2010). Inversely, increased ability for plastic changes at synaptic, cellular and higher levels of brain organization is indicative of an increased ability for plastic changes at the molecular level (Koehl 2015).

Hippocampus is perhaps the most representative brain region of a remarkably enhanced ability for plasticity at virtually all levels organization (Koehl 2015; Toda and Gage 2017; Rebola et al. 2017; Igarashi 2015; Sierra et al. 2015). However, the ability for plastic changes is not homogeneously distributed along the hippocampus formation, which in rodents extends from the septal region to the temporal lobe (for that the term “septotemporal” for its longitudinal axis). For instance, the ability for induced long-term synaptic potentiation, a widely accepted synaptic model for learning and memory (Takeuchi et al. 2014), and neurogenesis, a higher order plasticity phenomenon (Toda and Gage 2018), markedly differ along the dorsoventral axis of hippocampus, at last under some conditions (Papatheodoropoulos and Kostopoulos 2000; Kouvaros and Papatheodoropoulos 2016; Maruki et al. 2001; Colgin et al. 2004; Milior et al. 2016; Dubovyk and Manahan-Vaughan 2018a; Grigoryan and Segal 2013; Papaleonidopoulos and Papatheodoropoulos 2018; Schreurs et al. 2017; Moschovos and Papatheodoropoulos 2016; Banasr et al. 2006; Jayatissa et al. 2006; Boldrini et al. 2009). These, represent endogenous mechanistic aspects of the more general concept of functional segregation along the dorsoventral axis of hippocampus postulating that specific behavioral functions can be ascribed to distinct segments along the hippocampal formation, usually and shortly expressed by a dichotomy between cognition and emotionality related processes undertaken by the dorsal (DH) and the ventral hippocampus (VH) respectively (Bannerman et al. 2014; Strange et al. 2014; Gulyaeva 2018). The recently revealed striking diversification in patterns of gene expression along the hippocampus (Sotiriou et al. 2005; Pandis et al. 2006; Gusev et al. 2005; Leonardo et al. 2006; Thompson et al. 2008; Dong et al. 2009; Snyder et al. 2011; Cembrowski et al. 2016; Lee et al. 2017; Chawla et al. 2018; Floriou-Servou et al. 2018), may have profound functional implications for the hippocampal circuitry and the higher-level functional diversification along hippocampus. However, though some dorsal-ventral differences, for instance in synaptic and cell level have been successfully linked to molecular diversity along the hippocampus (Pandis et al. 2006; Dubovyk and Manahan-Vaughan 2018a; Snyder et al. 2011; Zhu et al. 2018; Floriou-Servou et al. 2018; Dubovyk and Manahan-Vaughan 2018b), the functional implications of diversified patterns of gene expression are largely unknown (Tushev and Schuman 2016).

A particularly important expression of the potentiality of a neural network for plastic changes is the ability to spontaneously generate physiological patterns of collective activities in the absence of an external triggering input (Tsukamoto-Yasui et al. 2007). Such an activity has been found in the hippocampus and consists of the so-called sharp wave-ripple complexes (Buzsaki 2015). Sharp waves (SPWs) are one of the dominant patterns of physiological collective activities of the hippocampus (Colgin 2016; Buzsaki 2015). Because SPWs are self organized in the denervated hippocampus (Buzsaki et al. 1987) and in isolated hippocampal preparations under normal in vitro conditions (Maier et al. 2003; Colgin et al. 2004; Kubota et al. 2003; Wu et al. 2002; Aivar et al. 2014; Mizunuma et al. 2014; Papatheodoropoulos and Kostopoulos 2002), they are considered an endogenous hippocampus activity pattern, which however occurs with vastly higher probability in VH than in DH preparations (Papatheodoropoulos and Kostopoulos 2002; Kouvaros and Papatheodoropoulos 2017; Colgin et al. 2004; Caliskan et al. 2016). Although the mechanisms that underlie the physiological triggering of SPW, either in vivo or in vitro, are not known, it has been previously postulated (Buzsaki 1989, 1996) and more recently indirectly corroborated by experimental evidence (Mizunuma et al. 2014) that SPWs may arise from plasticity-related synaptic reorganization. Furthermore, experimentally induced synaptic strengthening has been shown to either trigger or enhance SPWs (Buzsaki 1984; Behrens et al. 2005; Papatheodoropoulos 2010). For their part and by virtue of their recurrent nature, SPWs could significantly assist in molecular and synaptic reorganization that supports the formation of memory traces in hippocampus (Buzsaki 1989, 2015). However, whether SPWs induce plastic molecular changes is not yet known.

Spontaneous activity in neural networks is known to induce molecular and synaptic rearrangement with potential functional consequences (Tsukamoto-Yasui et al. 2007). For instance, recurring synchronized activity can induce changes in the expression of synaptic biochemical biomarkers in hippocampal slice cultures (Casanova et al. 2013) and convulsive seizures induce induction of immediate early genes (IEGs) expression (Link et al. 1995). Motivated by these observations we choose to study the expression of two early genes, c-fos and zif-268 and the expression of seven subunits of GABA_A_ receptor in DH and VH slices immediately after tissue preparation and at different times of tissue maintenance under normal in vitro conditions. GABA_A_ receptor plays a basic role in the generation of SPWs (Koniaris et al. 2011; Buzsaki 2015) and is also endowed with plastic properties induced by neuronal activity (Raol et al. 2006; Casanova et al. 2013) We find that *in vitro* development of SPWs in VH slices is associated with a region selective increase in the expression of specific GABA_A_ receptor subunits and IEGs. Possible multifold implications of the data are discussed.

## Methods

### Animals and hippocampal slice preparation

Male Wistar rats (RRID:RGD_10028) five to seven weeks old were used in the study. Rats were housed in the Laboratory of Experimental Animals of the Department of Medicine, University of Patras (licence No: EL-13-BIOexp-04). Animals were maintained under standard conditions of light-dark cycle (12hrs light –12hrs dark) and temperature (20-22 °C) with free access to food and water. Experiments were conducted in accordance with the European Communities Council Directive Guidelines for the care and use of Laboratory animals (2010/63/EU – European Commission) and they have been approved by the “Protocol Evaluation Committee” of the Department of Medicine of the University of Patras and the Directorate of Veterinary Services of the Achaia Prefecture of Western Greece Region (reg. number: 203173/1049, 22/08/2014). In particular, the brain was removed following decapitation under deep anaesthesia with diethyl-ether and placed in chilled standard artificial cerebrospinal fluid (ACSF, 2-4 °C) containing, in mM: 124 NaCl, 4 KCl, 2 CaCl_2_, 2 MgSO_4_, 26 NaHCO_3_, 1.25 NaH_2_ PO_4_ and 10 glucose; ACSF was equilibrated with 95% O_2_ and 5% CO2 gas mixture at a pH=7.4. Then the two hippocampi were excised free from the brain and sliced using a McIlwain tissue chopper. Transverse slices, 500-550 µm-thick, were prepared from the dorsal (septal) and the ventral (temporal) segment of the hippocampus extending between 1.0 mm and 3.0 mm from the dorsal and the ventral end, as previously described (Sotiriou et al. 2005). After their preparation, slices were immediately transferred to an interface type recording chamber where they were maintained continuously perfused with ACSF of the same composition as above described (rate ∼1.5 ml/min) and humidified with a mixed gas consisting of 95% O_2_ and 5% CO_2_ at a constant temperature of 30±0.5 °C. In autoradiography experiments, slices were instantly frozen in isopentane (−45 °C) and stored at –70 °C until use. The frozen tissue slices were mounted on the chucks of a Bright microtome (at –18 °C) and sections of 15 μm from the transverse slices were thaw-mounted onto microscope slides for the autoradiography experiment and on poly-l-lysine-coated microscope slides for the in situ hybridization procedure as described in following sections.

### Electrophysiological *in vitro* recordings

Recordings of field potentials started immediate following placement of hippocampal slices in the chamber and continued for a period of eight hours. Recordings were made from the CA1 stratum pyramidale using carbon fibers (7 μm-thick, Kation Scientific, Minneapolis, USA). Field potential recordings were acquired and amplified X500 and then filtered at 0.5 Hz–2 kHz using Neurolog amplifiers (Digitimer Limited, UK). Signal was digitized at 10 kHz and stored on a computer disk for off-line analysis using the CED 1401-plus interface and the Spike software (Cambridge Electronic Design, Cambridge, UK). Spontaneous field potentials consisted of synchronous events with a prominent positive phase of relatively small amplitude (25-250 μV), (Wu et al. 2002; Papatheodoropoulos and Kostopoulos 2002). Previous studies have demonstrated that these spontaneous events represents an *in vitro* model of hippocampal sharp waves (SPWs), (Maier et al. 2003; Kubota et al. 2003), and this activity is regularly observed in transverse slices prepared mainly from the ventral but not the dorsal hippocampus (Kouvaros and Papatheodoropoulos 2017; Papatheodoropoulos and Kostopoulos 2002; Colgin et al. 2004; Caliskan et al. 2016). Because the amplitude of SPWs was fairly small especially at the beginning of their emergence, thus, to make feasible their detection original records were down sampled (at 2 kHz) and low-pass filtered at 40 Hz before the detection procedure was applied. Then, individual SPWs were detected after setting a threshold at a level where all putative events were identified as verified by visual inspection as previously described (Giannopoulos and Papatheodoropoulos 2013). SPWs were quantified by their amplitude measured by the voltage difference between the positive peak and the baseline.

### Histochemistry

In order to compare the expression of mRNA of GABA_A_ receptor subunits and of early genes during *in vitro* maintenance of DH and VH slices the quantitative neurochemical method of *in situ* hybridization was used. Furthermore, in order to quantify GABA_A_ receptors a radioligand binding assay was used. Both methods were combined with autoradiography. These experiments were conducted in transverse hippocampal slices as described below in detail.

### In Situ Hybridization

Experiments of *in situ* hybridization for GABA_A_ receptor subunits mRNA expression were performed as previously described (Sotiriou et al. 2005). Specifically, the following synthetic oligonucleotides probes (MWG Biotechnology, UK) were used:

α1: 5’CCT GGC TAA GTT AGG GGT ATA GCT GGT TGC TGT AGG AGC ATA TGT 3’

α2: 5’AGG ATC TTT GGA AAG ATT CGG GGC GTA GTT GGC AAC GGC TAC AGC3’

α4: 5’CAA GTC GCC AGG CAC AGG ACG TGC AGG AGG GCG AGG CTG ACC CCG3’

α5: 5’TCC CCA GTC CCG CCT GGA AGC TGC TCC TTT GGG ATG TTT GGA GGA 3’

β1: 5’TGC CTG TCC AGC CCT CGT CCG AAG CCC TCA CGG CTG CTC AGT GGT 3’

β2: 5’ACT GTT TGA AGA GGA ATC TAG TCC TTG CTT CTC ATG GGA GGC TGG 3’

β3: 5’CTG TCT CCC ATG TAC CGC CCA TGC CCT TCC TTG GGC ATG CTC TGT 3’

γ2: 5’CAT TTG GAT CGT TGC TGA TCT GGG ACG GAT ATC AAT GGT AGG GGC 3’

δ: 5’GAG GGA GAA GAG GAC AAT GGC GTT CCT CAC GTC CAT CTC TGC CCT 3’

The probes were 3’ end-labeled, using 30:1 molar ratio of isotope [^35^S] dATP (1250Ci/mmol, NEN) to oligonoucleotide and terminal deoxynucleotidyl transferase (TDT, Roche). Non incorporated nucleotide was removed by Biospin 6 chromatography columns (Bio-rad). Before the hybridization procedure sections were transferred from –20 °C to room temperature and allowed to air dry. Then sections were fixed mildly in 4% paraformaldehyde for 5 minutes, washed for 1 minute in phosphate buffer saline (PBS), dehydrated (in 70%, 95% and 100% ethanol, for 5 minutes at each solution) at room temperature and allowed to air dry. The slides were then incubated for 18 hours at 42 °C in 90 μl of hybridization buffer containing (∼2 × 10^5^ cpm) of labeled probe. Nonspecific signal was determined by addition of 100-fold excess of unlabelled probe to the hybridization solution on some slides. Hybridization buffer consists of 50% v/v formamide, 4xSSC (1xSSC is 0.15 M sodium chloride and 0.015 M sodium citrate) and 10% w/v dextran sulfate. Following hybridization, sections were washed several times in 1xSSC at 60 °C and room temperature, dehydrated, allowed to air dry and exposed to X-ray film (Biomax MR Kodak) for 4-6 weeks at room temperature.

Experiments of *in situ* hybridization for early genes mRNA expression were performed as previously described (Dassesse et al. 1999). In particularly, as probes, DNA oligonucleotides with a complementary base sequence with the portion of the mRNA expressing either the early genes *c-fos* and zif-268 were used, and the base sequence was 3’CGG TGC AGC CAT CTT ATT CCG TTC CCT TCG GAT TCT 5’ and 5’GCG GCG AAT CGC GGC GGC TCC CCA AGT TCT GCG CGC TGG GAT CTC 3’ for *c-fos* and zif-268 respectively.

Each probe for both the seven GABA_A_ receptor subunits and the two early genes was diluted at a concentration of 3 pmoles/ml and labeled with ^35^S-ATP at the 3’ end with the 3’ terminal transferase in order to have a specific activity of ~ 6X10^5^ cpm/µl. Non incorporated nucleotides were removed by exclusion chromatography (Sephadex G-50). The hybridization solution was placed on the sections, which were immediately covered with parafilm, placed in plastic plug and incubated at 42 °C for 18 hours. The sections were then immersed in 1X SSC at room temperature, washed in 1X SSC for 20 minutes at 60 °C, then immersed in 1X SSC at room temperature, rinsed in 0.1X SSC for 3 minutes at room temperature, dehydrated in increasing concentrations of ethanol (70%, 95%, 100%) and allowed to dry. Non-specific binding was determined in the presence of a 100-fold excess unlabelled probe.

### Receptor autoradiography

Autoradiography experiments were performed as previously described (Titulaer et al. 1994). Shortly, on the day of the experiment the sections underwent a pre-wash for 10 minutes (twice) in 50 mM Tris-citrate buffer at 4 °C, pH 7.2. The sections were then incubated for 40 minutes at 4 °C in 50 mM Tris-citrate buffer, pH 7.2 containing 60nM [^3^H]muscimol (s.a 20 Ci/mmol, Amersham, Netherlands). Non-specific binding was determined as binding in the presence of 0.1 mM GABA in adjacent sections, which represented less than 10% of the total binding. After incubation, the sections were washed in cold Tris-citrate 50 mM buffer, pH 7.2 for 1 minute (3 times) followed by a dip into ice-cold distilled water and finally dried at the stream of cold air. Dried sections were exposed to tritium-sensitive film ([^3^H]-Hyperfilm, Amersham) along with tritium standards (Amersham) and stored in an X-ray film cassette for ~4 weeks at 4 °C.

### Quantitative image analysis

The *in situ* hybridization autoradiographs were developed with Kodak GBX developer and fixed with Kodak GBX fixer at 22 °C. Hippocampal fields (i.e. CA1, CA3 and Dentate Gyrus, DG) were quantified according to Paxinos and coll. atlas (Paxinos et al. 2009). Quantification analysis of the resulting autoradiographic images was performed by using the MCID image analysis system (M1, Imaging Research, Canada). Optical density measurements from each area were defined as relative optical density (ROD) units. Measurements were made in 6-8 sections from each hippocampal segment (DH, VH) obtained from eight to ten rats. Non-specific signal represented less than 5% of the total signal and therefore was not evaluated.

Receptor binding autoradiographs were developed with Kodak D-19 developer and fixed with Kodak Acidofix at 18 °C. Quantification analysis of autoradiographs was performed as above described. Optical density measurements from each field in DH and VH were converted into fmol [^3^H]muscimol per mg tissue equivalent, according to the calibration curve obtained from the tritium standards. Measurements were performed in six sections of each hippocampal segment (DH, VH) obtained from each rat in a total of 10 rats. **Statistics**

The parametric independent t-test and the one-way analysis of variance test (ANOVA, with Dunnet post-hoc test) were used to evaluate statistically significant differences. Statistical significance was assessed at *p*<0.05. Detection of statistically significant differences was made using the number of animals. The values of variables throughout the text represent the means±S.E.M.

## Results

### *In vitro* development of sharp waves (SPWs) in ventral hippocampal slices

Field recordings started immediately after placing hippocampal slices in the recording chamber and continued over the next eight hours. We observed that spontaneous field activity of very small amplitude (<10 μV) could be observed from VH but not DH slices soon after tissue was placed in the recording chamber (i.e. during the first half an hour of tissue maintenance). However, a detection and therefore measurement of SPWs, without the risk that physiological activity being contaminated with noise was technically feasible starting at thirty minutes of slice maintenance (Fig. 1). At this time spontaneous activity was detected in the great majority of VH slices (n=56/67) but in only two DH slices (n=2/52). This is in keeping with previous studies that have shown that ventral but not dorsal transverse hippocampal slices have an increased ability to spontaneously generate SPWs under normal *in vitro* conditions (Papatheodoropoulos and Kostopoulos 2002; Kouvaros and Papatheodoropoulos 2017; Colgin et al. 2004; Caliskan et al. 2016). Actually, SPWs can be generated in a small fraction of DH slices as shown by assessment of a large population of slices in a recent study (Kouvaros and Papatheodoropoulos 2017). However, in the limited population of DH slices used in the present study we could not reliably and undeniably detect SPWs. As previously demonstrated, this activity represents an *in vitro* analogue of sharp waves (SPWs) recorded from the hippocampus of behaving animals that represents a very accurately structured physiological activity pattern that is generated endogenously in the hippocampus and plays a crucial role in the process of memory consolidation, among other functions (Buzsaki 2015). SPWs, as measured by their amplitude, developed gradually during the first 3-4 hours of tissue maintenance, thereafter reaching a plateau (Fig. 1B, C). In particular, the amplitude of SPWs significantly increased from fifteen minutes to four hours of tissue maintenance *in vitro* (ANOVA at 15 min - 4 hours, *p* < 0.005), while it presented no further significant change afterwards (ANOVA at four hours-eight hours, *p* > 0.05). A decline in amplitude of SPWs was observed at late stages of tissue maintenance (8 hours) but it was not statistically significant (ANOVA post-hoc at 15 min-8 hours and independent t-test between 8 hours and 3.5, 4, 4.5, 5 hours, *p* > 0.05). To our best knowledge, this is the first time that the gradual development of *in vitro* SPWs under normal conditions is quantitatively described and characterized. Previous studies have shown that spontaneous activity can trigger rearrangement of molecular and/or cellular components influencing neuronal network functioning (Guzowski et al. 2001; Miyashita et al. 2008; Tsukamoto-Yasui et al. 2007). We therefore asked whether the occurrence of SPWs could be accompanied by plastic changes at the molecular level. Thus, we measured the expression of mRNA coding for the various subunits of GABA_A_ receptor and the mRNA expression for two immediately early genes, *c-fos* and zif-268, as described in the following sections.

**Figure 1.**
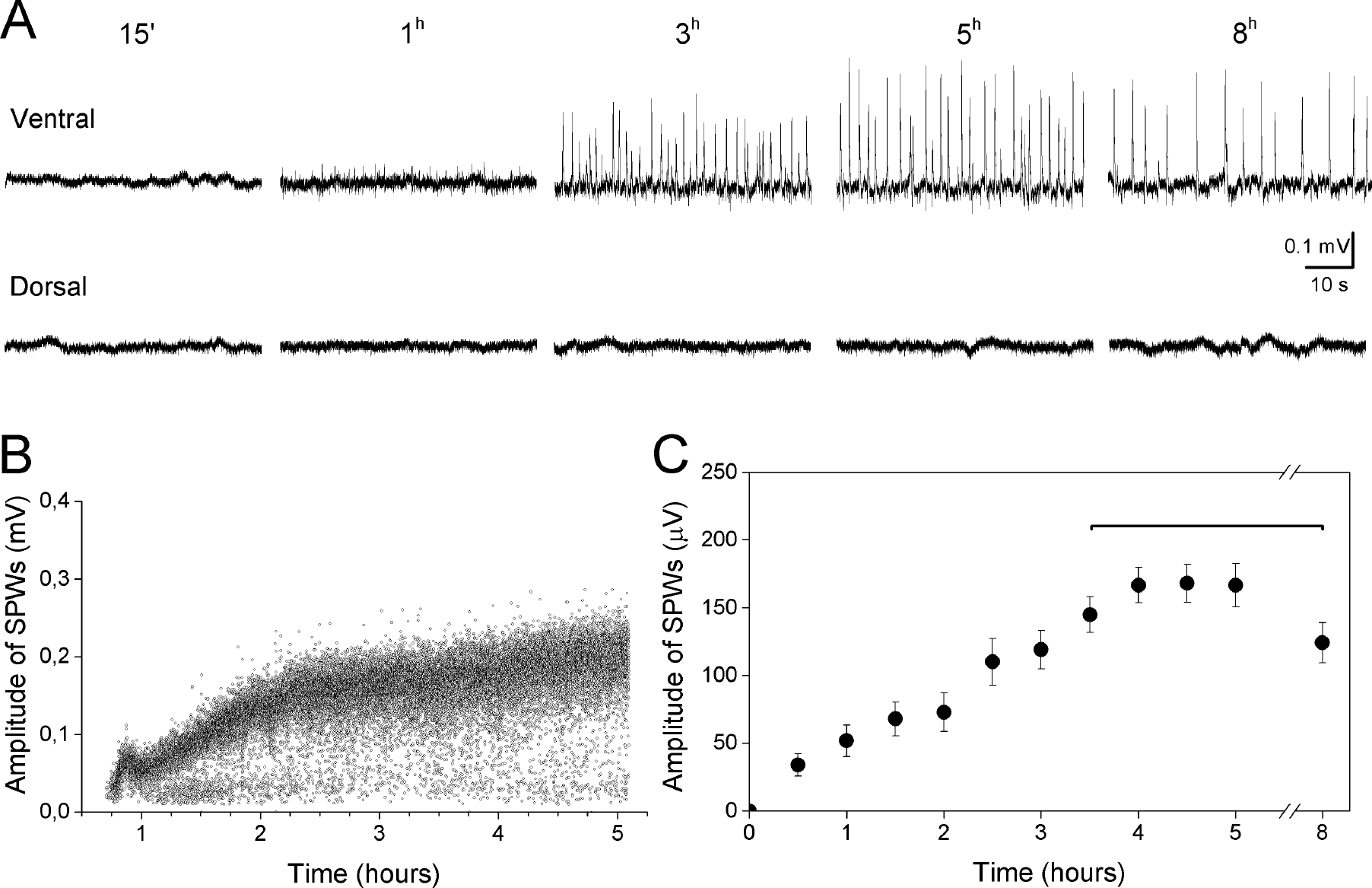
*In vitro* development of spontaneous sharp waves (SPWs) in ventral but not dorsal hippocampal slices. **A**. Continuous field potential recordings from the CA1 stratum pyramidale obtained at four representative time periods during slice maintenance in the recording chamber. **B**. Instantaneous time histogram of SPWs’ amplitude prepared from a continuous record five hours long obtained from a VH slice illustrating the time course of development of SPWs. **C**. Cumulative results on *in vitro* development of SPWs. Data were obtained from twenty five VH slices.

### GABA_A_ receptor reorganization is increased in the VH slices

The GABA_A_ receptor is formed by a combination of five subunits that belong to different families (Sigel and Steinmann 2012). GABA_A_ receptor is crucially involved in the generation of SPWs and some GABA_A_ receptor subtypes appear to play more important roles in SPWs compared with other subtypes; for a review, see (Buzsaki 2015). For instance, selective pharmacological activation of α1- or restricting α5-containing GABA_A_ receptor subtypes, which represent the most abundant GABA_A_ receptor subtypes in the hippocampus (Nusser et al. 1996; Mortensen et al. 2012) considerably facilitate SPWs’ generation (Koniaris et al. 2011). Furthermore, the α1-containing GABA_A_ receptor predominately mediates the actions of parvalbumin-expressing basket cells (Somogyi et al. 1996; Thomson et al. 2000; Klausberger et al. 2002), which play a critical role in SPWs (Somogyi et al. 2014). On the other hand, α2-containing GABA_A_ receptors mediate the actions of cholecystokinin-expressing cells (Nusser et al. 1996; Nyiri et al. 2001) the activation of which is invariant during SPWs (Somogyi et al. 2014). It is also worth noting that the α1-containing and α2-containing GABA_A_ receptors also preferentially contain the β2 and the β1 subunit respectively (Somogyi et al. 1996; Nusser et al. 1996). We therefore choose to examine the expression of mRNA coding for the following seven different subunits of GABA_A_ receptor: α1, α2, α5, β1, β2, β3 and γ2 subunit. The expression of mRNA was assessed at six time points during hippocampal tissue preparation and maintenance *in vitro*. Specifically, subunit mRNA expression was measured in dorsal and ventral hippocampal slices immediately following their preparation (time zero) and at 15 minutes, 30 minutes, 1 hour, 3 hours, 5 hours and 8 hours of slice maintenance in the chamber for electrophysiological recording.

We found significant and consistent increases in mRNA expression for particular GABA_A_ receptor subunits in DH and VH in the CA1 field; however no consistent pattern of mRNA changes was observed in dentate gyrus and CA3 field in either DH or VH slices (Fig. 2 and Fig. 3). Thus, highly increased mRNA levels were found for α1, β2, and γ2 subunits in the CA1 field of VH slices during development of spontaneous SPWs (Fig. 3). Specifically, the mean increase in mRNA expression during the period from one to eight hours *in vitro* was 21.76±2.83%, 34.54±3.51% and 16.43±1.48% for α1, β2, and γ2 subunits in VH slices respectively (independent t-test for each subunit, *p*<0.05). In the CA1 field of DH slices a significant mean increase in mRNA expression was observed for subunits α2 (16.0±1.81%) and α5 (10.74±0.72), (independent t-test for each subunit, *p*<0.05). In all these cases significant increase in mRNA levels was found starting at one hour after slice placement in the recording chamber (for additional statistics see Fig. 3). No significant changes in mRNA expression was observed for subunits α1, β2, γ2, β1, and β3 in DH slices and for subunits α2, α5, β1, and β3 in VH slices. It should also be noted that the baseline expression of mRNA (i.e. in naive slices) for α1, β2, and γ2 subunits was significantly higher in DH than VH slices (independent t-test between DH and VH for each subunit, *p*<0.05), while the baseline mRNA expression for α2, α5 and β1 subunits was higher in the VH compared with the DH (independent t-test, *p*<0.05), corroborating thus previous observations (Sotiriou et al. 2005). It is interesting that time-dependent increase in mRNA expression in VH slices was observed for those subunits whose baseline expression was reduced in VH (i.e. α1, β2, and γ2). Correspondingly, increase in mRNA expression in DH slices maintained in vitro was observed for α2 and α5 subunits the level of which is lower in DH than VH naive slices. Furthermore, during eight-hour slice maintenance *in vitro* the mRNA levels for subunits β2, γ2 and α5 (but not α1 and α2) became similar in DH and VH (independent t-test between the two segments for each subunit, *p*>0.05).

**Figure 2.**
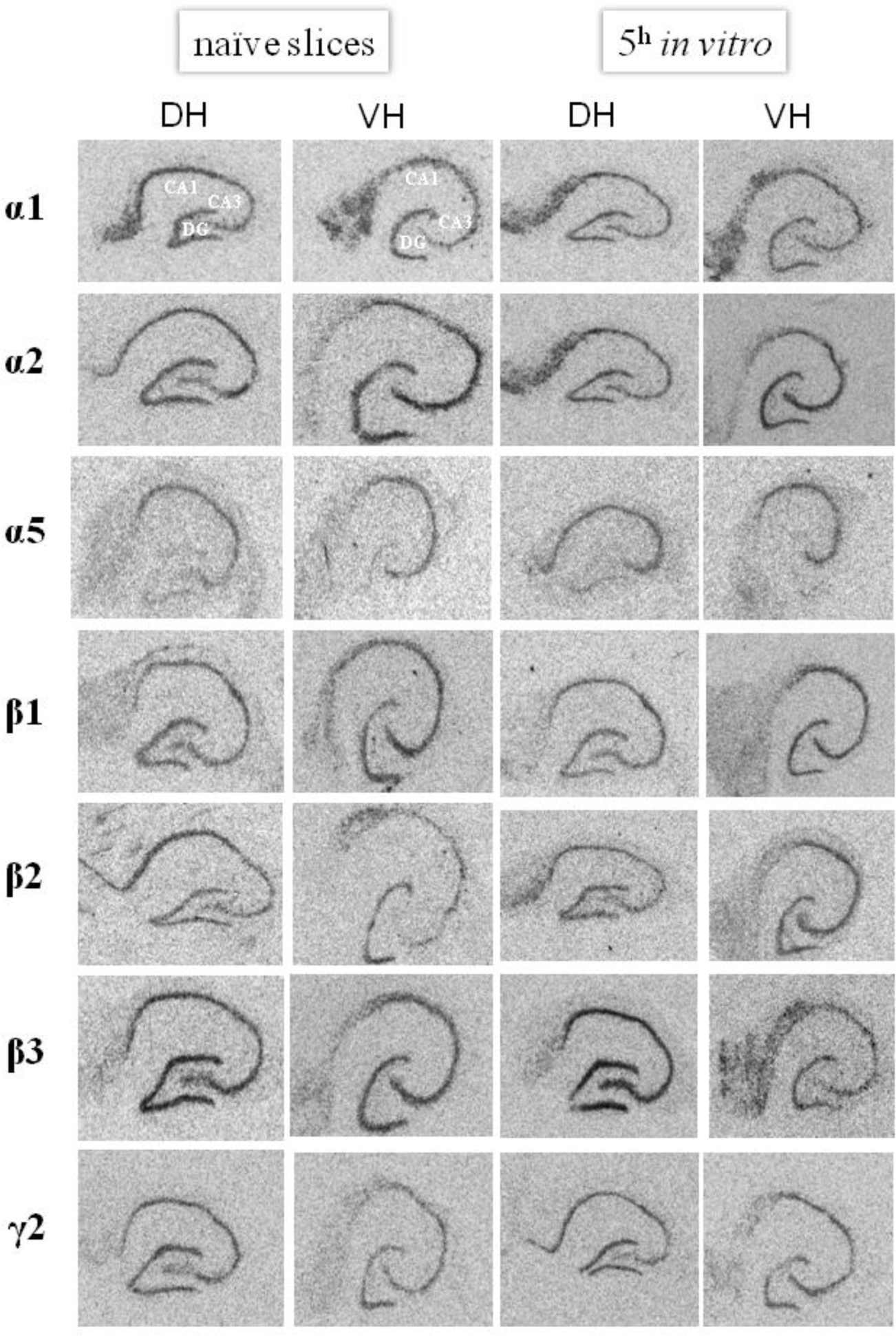
Examples of *in situ* hybridization images obtained by film autoradiography illustrating mRNA expression for seven GABA_A_ receptor subunits, obtained from dorsal (DH) and ventral (VH) hippocampal slices immediately following their preparation (naive slices) and 5 hours of their maintenance *in vitro* (5h *in vitro*).

**Figure 3.**
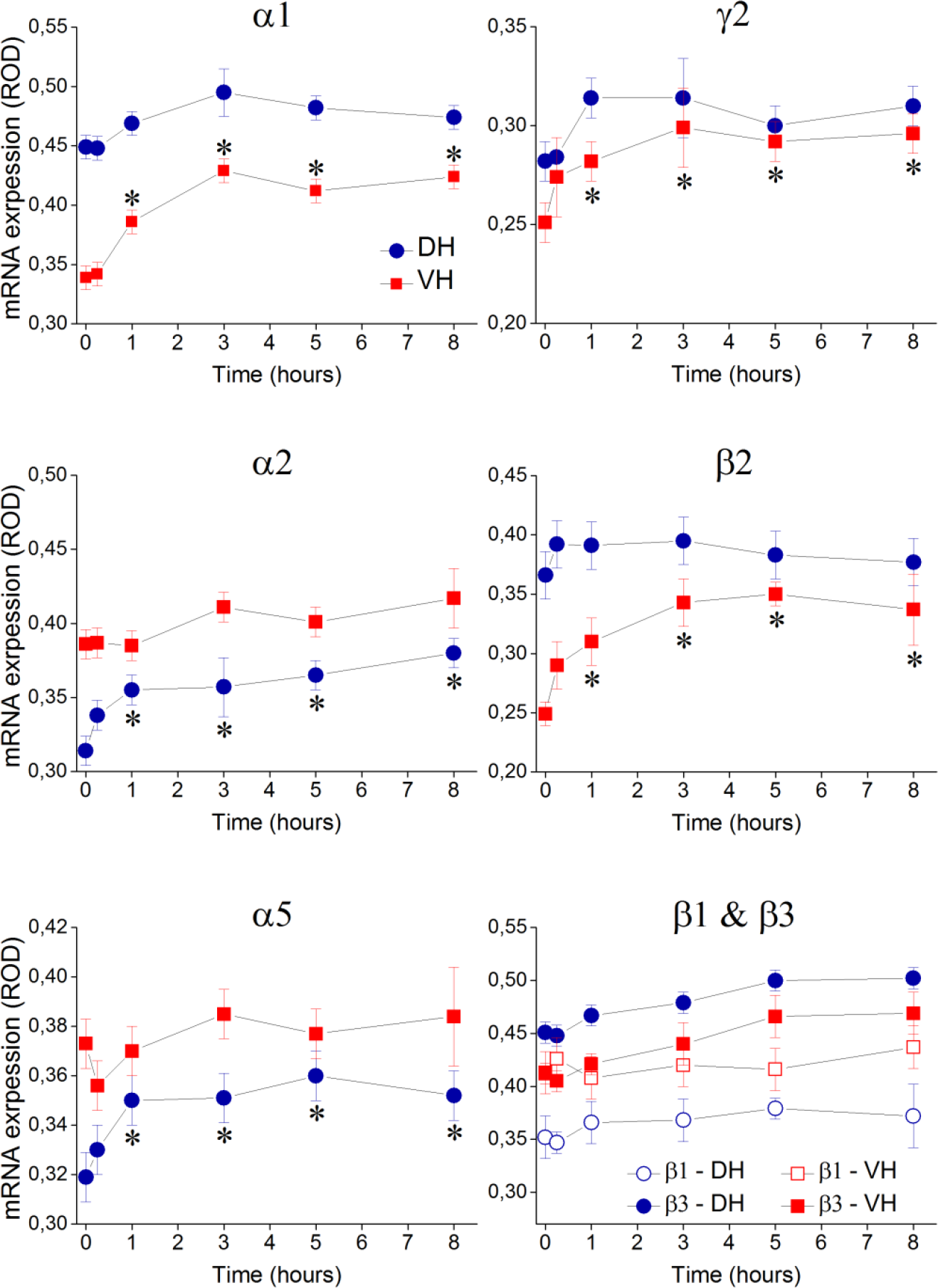
Time profiles of expression of mRNA coding for seven GABA_A_ receptor subunits in the CA1 field of DH and VH slices maintained *in vitro*. Profiles extent from time zero (measurements obtained immediately after slice preparation) until eight hours of tissue maintenance *in vitro*. Cumulative data obtained from ten rats are shown. Asterisks indicate statistically significant differences in mRNA expression during slice maintenance in the recording chamber with respect to expression at time zero (ANOVA, *p*<0.05). Note that significant changes in mRNA expression in the VH occur for those subunits that are expected to form functional GABA_A_ receptor subtype (α1/β2/γ2).

Regarding the mRNA expression in hippocampal fields CA3 and dentate gyrus, we observed some significant changes that were in partial consistency with the changes in CA1 field. In particular, in the ventral CA3 field significant increase in expression occurred in mRNAs coding for α1 and β2 subunits while in the dorsal CA3 field increased mRNA levels were observed for α5 and β3 subunits (Table 1). Surprisingly, in the dentate gyrus we observed reductions in the mRNA levels coding for some subunits in the DH (α1 and β3 subunits) and the VH (γ2 subunit), (Table 2). These results showed that *in vitro* maintenance of hippocampal slices is accompanied by region-specific molecular reorganization in GABA_A_ receptor subunit expression. Importantly, the molecular changes are more robust and selective in the region of VH slices where spontaneous network activity is organized (CA1) compared with other hippocampal fields and also compared with “silent” DH slices.

**Table 1.** Time profiles of mRNA expression (ROD units) for seven GABA_A_ receptor subunits in the **CA3** field of DH and VH slices maintained *in vitro*.

| Time *in vitro*: |  | 0 (naive) | 15 min | 1 h | 3 h | 5 h | 8 h |
| --- | --- | --- | --- | --- | --- | --- | --- |
| $\alpha 1$ | DH | 0,388±0,01 | 0,368±0,02 | 0,402±0,01 | 0,426±0,01 | 0,401±0,01 | 0,414±0,01 |
|  | VH | 0,338±0,01 | 0,348±0,01 | 0,354±0,01 | 0,392±0,01* | 0,378±0,01* | 0,397±0,01* |
| $\alpha 2$ | DH | 0,347±0,01 | 0,374±0,01 | 0,390±0,02 | 0,393±0,02 | 0,423±0,02 | 0,414±0,01 |
|  | VH | 0,428±0,01 | 0,423±0,02 | 0,420±0,03 | 0,437±0,01 | 0,461±0,02 | 0,446±0,01 |
| $\alpha 5$ | DH | 0,345±0,01 | 0,336±0,01 | 0,359±0,01 | 0,389±0,01* | 0,394±0,01* | 0,373±0,01* |
|  | VH | 0,378±0,01 | 0,359±0,01 | 0,382±0,01 | 0,405±0,01 | 0,392±0,01 | 0,404±0,01 |
| $\beta 1$ | DH | 0,349±0,02 | 0,348±0,02 | 0,362±0,02 | 0,357±0,02 | 0,384±0,01 | 0,377±0,01 |
|  | VH | 0,407±0,02 | 0,411±0,02 | 0,401±0,02 | 0,407±0,03 | 0,415±0,02 | 0,408±0,02 |
| $\beta 2$ | DH | 0,347±0,02 | 0,323±0,02 | 0,309±0,02 | 0,337±0,02 | 0,319±0,02 | 0,324±0,02 |
|  | VH | 0,295±0,02 | 0,299±0,02 | 0,304±0,02 | 0,386±0,06* | 0,319±0,02* | 0,343±0,02* |
| $\beta 3$ | DH | 0,424±0,01 | 0,407±0,01 | 0,410±0,01 | 0,476±0,01* | 0,468±0,01* | 0,477±0,01* |
|  | VH | 0,409±0,01 | 0,376±0,01 | 0,414±0,02 | 0,439±0,02 | 0,438±0,02 | 0,432±0,01 |
| $\gamma 2$ | DH | 0,274±0,01 | 0,258±0,01 | 0,289±0,01 | 0,309±0,01 | 0,290±0,01 | 0,298±0,01 |
|  | VH | 0,289±0,01 | 0,285±0,01 | 0,281±0,01 | 0,293±0,01 | 0,278±0,01 | 0,307±0,01 |

\*Comparison of value at indicated time point with value at time=0; independent t-test at $p<0.05$ .

Table 1. Time profiles of mRNA expression (ROD units) for seven GABA<sub>A</sub> receptor subunits in the **DG** field of DH and VH slices maintained *in vitro*.
| Time *in vitro*: |  | 0 (naive) | 15 min | 1 h | 3 h | 5 h | 8 h |
| --- | --- | --- | --- | --- | --- | --- | --- |
| $\alpha 1$ | DH | 0,416±0,01 | 0,381±0,01 | 0,380±0,02 | 0,398±0,01 | 0,364±0,01* | 0,356±0,01* |
|  | VH | 0,395±0,01 | 0,356±0,02 | 0,347±0,01 | 0,386±0,01 | 0,361±0,01 | 0,351±0,01 |
| $\alpha 2$ | DH | 0,380±0,01 | 0,362±0,01 | 0,379±0,01 | 0,365±0,01 | 0,396±0,02 | 0,352±0,02 |
|  | VH | 0,379±0,02 | 0,425±0,01 | 0,393±0,01 | 0,407±0,01 | 0,415±0,01 | 0,382±0,01 |
| $\beta 1$ | DH | 0,397±0,02 | 0,359±0,02 | 0,379±0,01 | 0,366±0,01 | 0,356±0,02 | 0,343±0,02 |
|  | VH | 0,402±0,03 | 0,394±0,02 | 0,380±0,02 | 0,402±0,02 | 0,384±0,01 | 0,366±0,01 |
| $\beta 2$ | DH | 0,350±0,02 | 0,333±0,02 | 0,330±0,02 | 0,324±0,02 | 0,309±0,02 | 0,320±0,03 |
|  | VH | 0,342±0,02 | 0,335±0,02 | 0,304±0,02 | 0,317±0,02 | 0,325±0,02 | 0,304±0,02 |
| $\beta 3$ | DH | 0,582±0,01 | 0,526±0,01 | 0,538±0,02 | 0,531±0,01 | 0,497±0,01 | 0,510±0,01 |
|  | VH | 0,467±0,01 | 0,400±0,01 | 0,426±0,01 | 0,438±0,01 | 0,441±0,02 | 0,469±0,01 |
| $\gamma 2$ | DH | 0,350±0,01 | 0,328±0,01 | 0,318±0,01 | 0,333±0,01 | 0,290±0,01* | 0,285±0,01* |
|  | VH | 0,309±0,01 | 0,301±0,01 | 0,302±0,01 | 0,319±0,01 | 0,286±0,01 | 0,285±0,01 |

\*Comparison of value at indicated time point with value at time=0; independent t-test at $p<0.05$ .

### GABA_A_ receptor reorganization is accompanied by an increased receptor number in the VH but not the DH

The results on receptor subunit mRNA expression showed a significant time-dependent increase in the mRNA expression coding for the α1, β2 and γ2 subunits in the VH and α2 and α5 subunits in the DH. It has been previously shown that the α1 subunit-containing is the most abundant GABA_A_ receptor subtype in the brain while the α2 subunit-containing GABA_A_ receptors represent a minority (Benke et al. 2004; Engin et al. 2018). Furthermore, GABA_A_ receptors composed of α1, β2 and γ2 subunits are fully functional receptors (Nusser et al. 1996; Benke et al. 2004). We therefore hypothesized that the increase in mRNA expression for these subunits in the VH could lead to an increase in the number of functional GABA_A_ receptors specifically in this segment of hippocampus. We examined the expression of GABA_A_ receptors by measuring [^3^H]muscimol binding in slices maintained eight hours in the recording chamber (Fig. 4). As shown in figure 5, [^3^H]muscimol binding significantly increased in all fields of VH slices (independent t-test for each field and layer, *p*<0.05). On the contrary, [^3^H]muscimol binding in DH slices remained in control levels (independent t-test for each field and layer, *p*>0.05). These results suggested that the number of functional GABA_A_ receptors increased in VH but not in DH slices during their maintenance *in vitro*.

**Figure 4.**
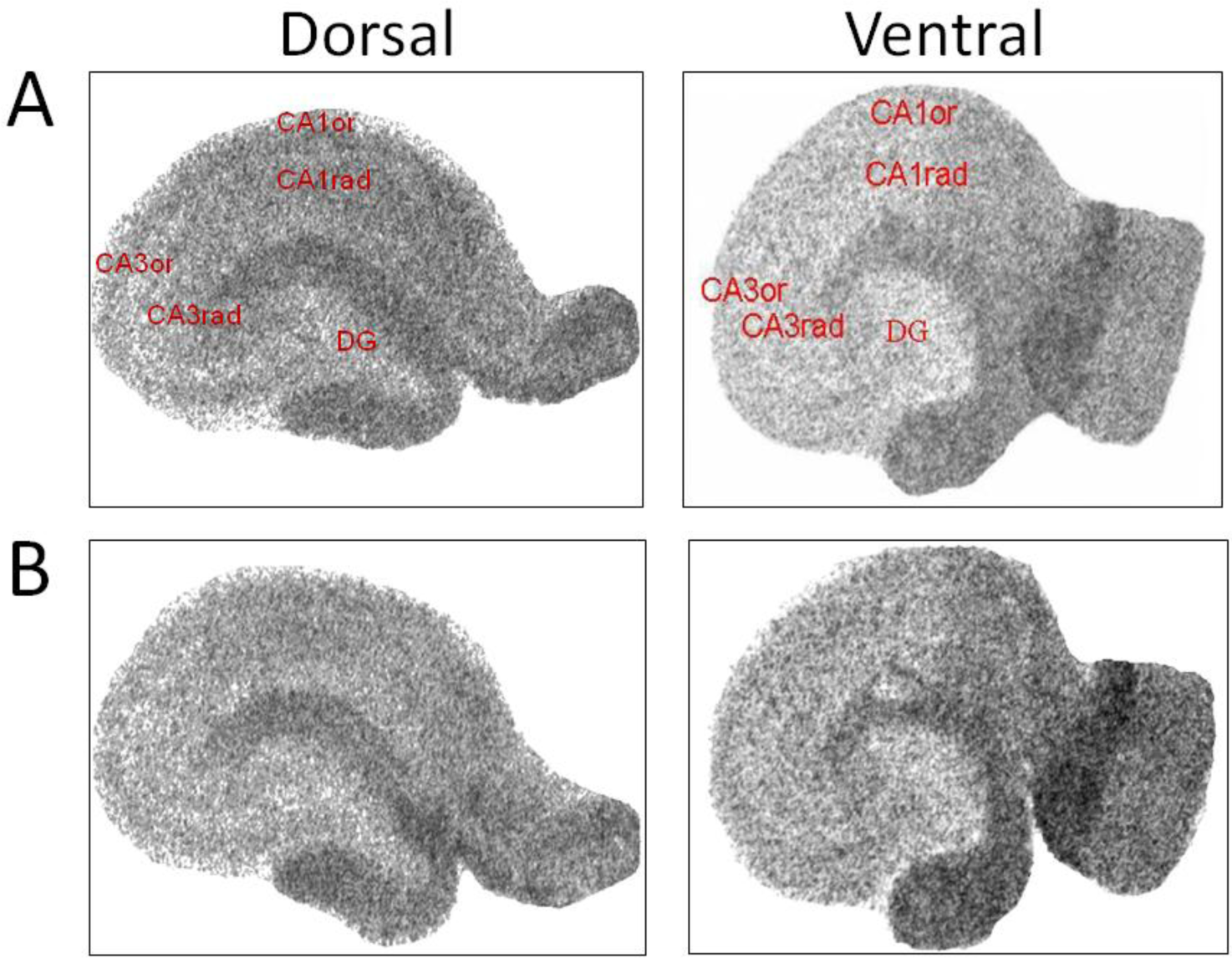
Examples of quantitative autoradiograms of ^[3]^muscimol binding obtained from naive dorsal and ventral slices (A) and from slices maintained eight hours in the recording chamber. The three hippocampal subfields (DG, CA3 and CA1) and the two layers of CA fields, i.e. stratum oriens (CA3or, CA1or) and stratum radiatum (CA3rad, CA1rad), used for [^3^H]muscimol binding measurements are indicated in pictures shown in A.

**Figure 5.**
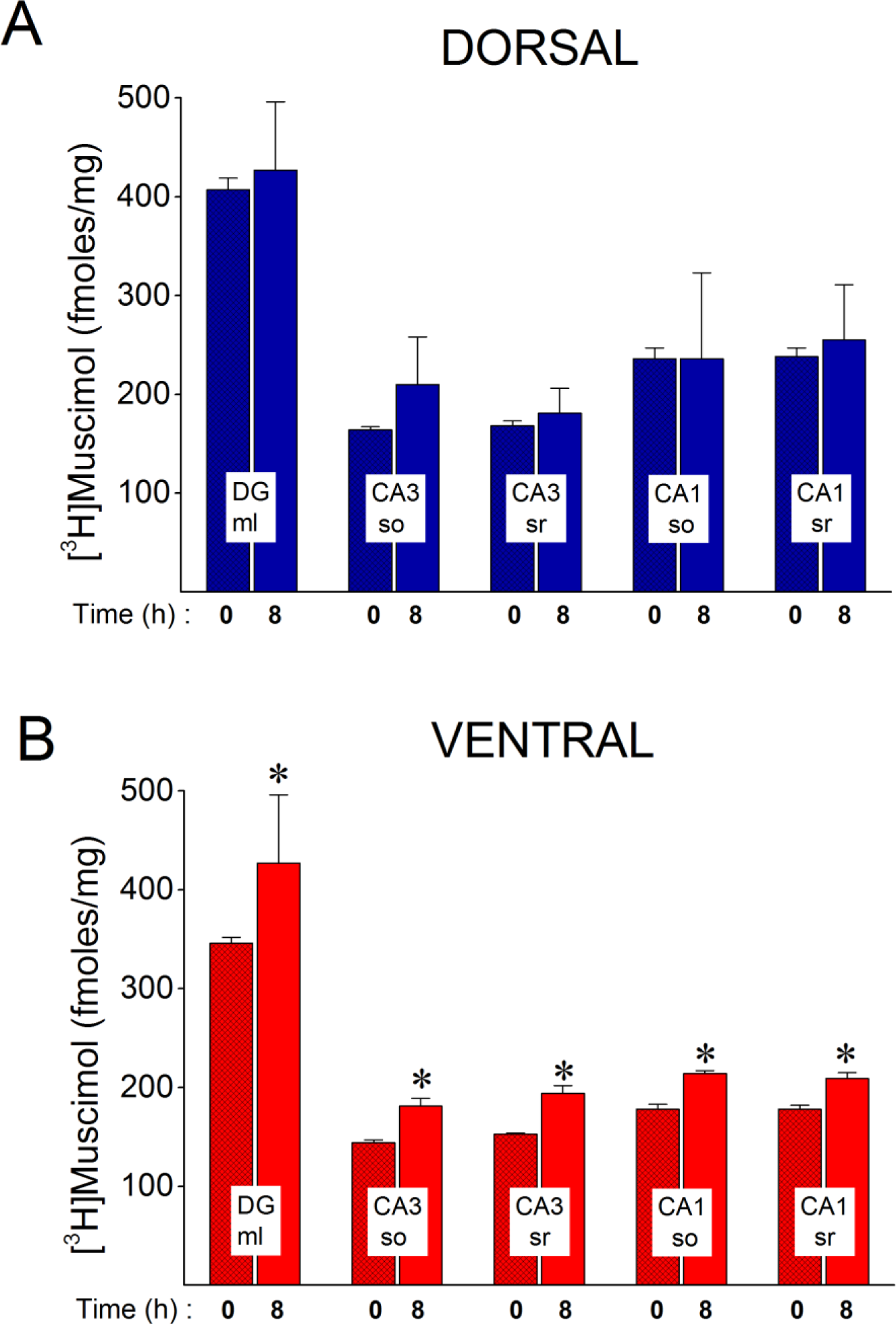
[^3^H]muscimol binding significantly increases during *in vitro* maintenance of VH (B) but not DH slices (A). Binding was measured in five hippocampal layers: molecular layer of dentate gyrus (DG ml), stratum oriens and stratum radiatum of CA3 and CA1 fields (CA3 so, CA3 sr, CA1 so, CA1 sr respectively). Time zero marks the time of slice preparation. Asterisks indicate statistically significant differences in [^3^H]muscimol binding measured eight hours of slice maintenance in the recording chamber compared with freshly prepared slices (independent t-test, *p*<0.05).

**Figure 6.**
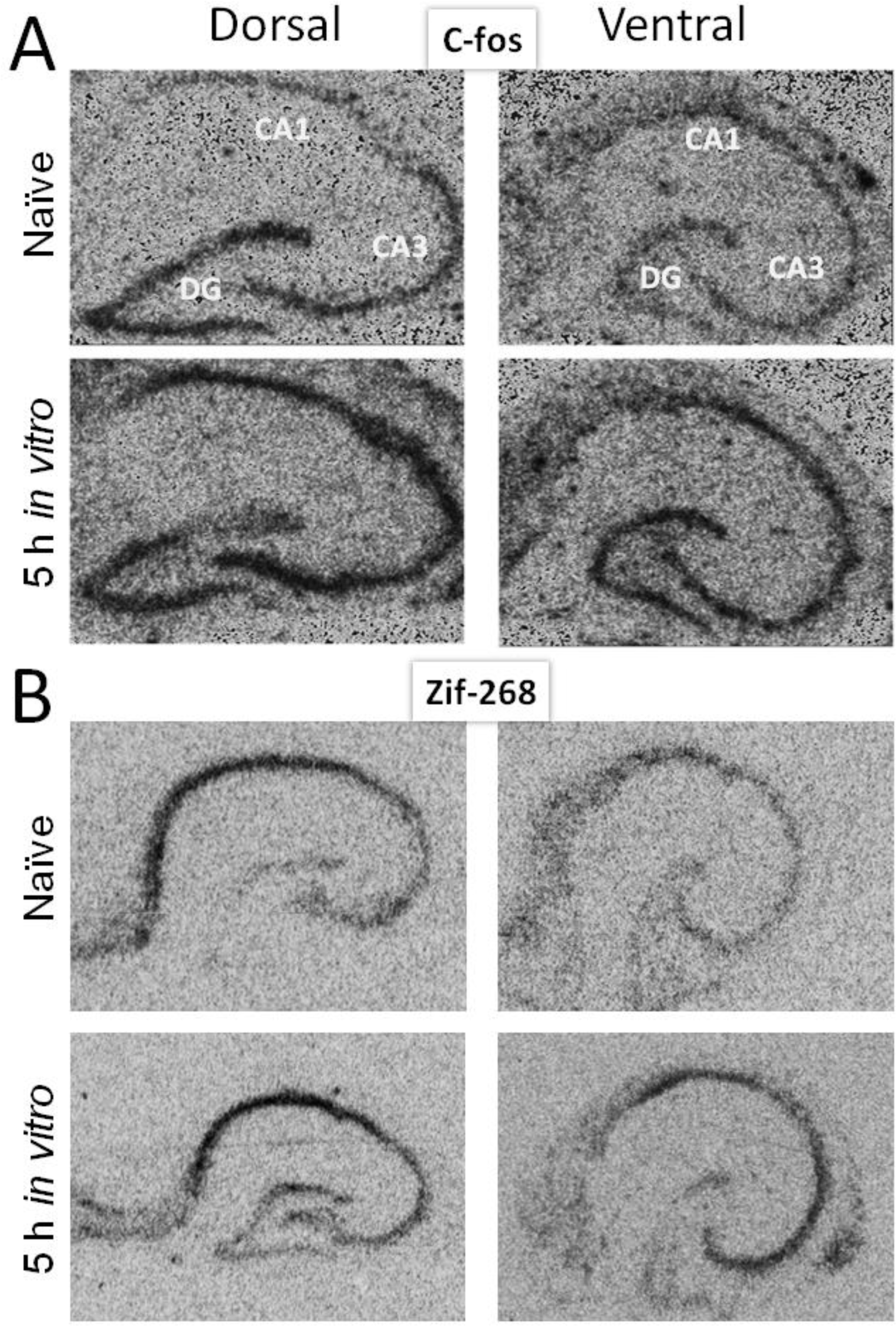
Examples of *in situ* hybridization autoradiographic images illustrating mRNA expression of *c-fos* (A) and zif-268 (B) in dorsal and ventral hippocampal slices immediately after their preparation (Naive) and after five hours of maintenance in the electrophysiological recording chamber (5 h *in vitro*).

### Increased expression in immediately early gene expression accompanies *in vitro* development of SPWs in the VH

Expression of early genes can occur following a number of triggering factors including electrical activity of neurons (Miyashita et al. 2008). In order to examine whether the occurrence of spontaneous activity can induce the expression of early genes selectively in VH slices we examined the levels of mRNAs in naive slices and in slices maintained for five hours in the recording chamber, i.e. when spontaneous activity was fully developed in VH slices. We found that both *c-fos* and zif-268 were expressed in all hippocampal fields of naive slices. Furthermore the expression was similar in DH and VH (independent t-test, *p*>00.5), (Fig. 7). During the *in vitro* tissue maintenance the expression of both genes significantly increased in DG and CA3 of both hippocampal segments (independent t-test, *p*<0.05). Strikingly, however, the expression of both *c-fos* and zif-268 in the CA1 field significantly increased in VH but not in DH slices (Fig. 7). Specifically, the expression of *c-fos* significantly increased in either hippocampal segment by 23% in DG (independent t-test in each hippocampal segment, *p*<0.05) and 24-26% in CA3 (independent t-test in each hippocampal segment, *p*<0.05). Similarly, the expression of zif-268 increased in both hippocampal segments by 50% in DG and 33-48% in CA3 (independent t-test in each hippocampal segment and field, *p*<0.05). However, the expression of *c-fos* and zif-268 in the CA1 field significantly increased in the VH (by 17% and 60% respectively, independent t-test, *p*<0.05) but not in DH (independent t-test, *p*>0.05).

**Figure 7.**
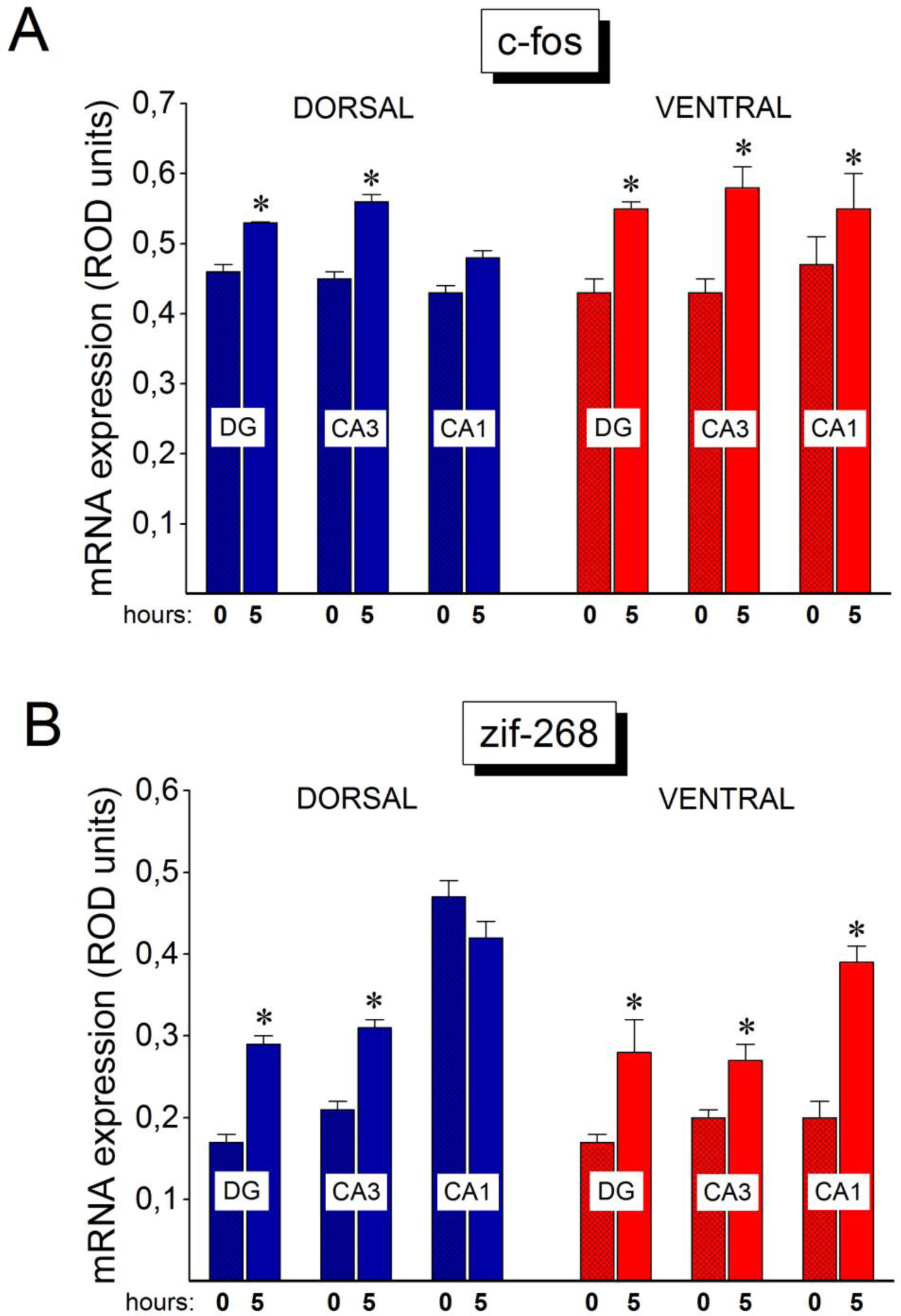
The expression of early genes significantly increases in the CA1 hippocampal field of VH but not DH during the *in vitro* tissue maintenance. Early genes expression was measured in the three hippocampal fields, DG, CA3 and CA1 immediately after slice preparation (marked by time zero) and at five hours of tissue maintenance in the recording chamber. Asterisks indicate statistically significant differences in mRNA expression measured eight hours of slice maintenance in the recording chamber compared with freshly prepared slices (independent t-test, *p*<0.05). The expression of early genes is significantly increased in DG and CA3 of both hippocampal segments, however, the expression increases in the CA1 field of only VH. Note that the increase in early genes expression parallels the *in vitro* development of SPWs in VH slices.

## Discussion

The main findings of the present study are as follows: a) In vitro maintenance of hippocampal slices is accompanied by significant changes in tissue molecular organization and the development of a physiological activity pattern in VH but not DH slices; b) The gradual and spontaneous in vitro development of SPWs in VH slices is associated with a concerted increase in the expression of mRNA coding for specific GABA_A_ receptor subunits (α1, β2, γ2) and leads to increase GABA_A_ receptor quantity. Instead, an increase in the mRNA expression for some GABA_A_ receptor subunits (α2, α5) in DH slices is not followed by quantitative receptor changes; c) Full development of SPWs in CA1 field of VH slices is followed by robust expression of IEG (c-fos, zif-268) in the same hippocampal field. In the DH slices, which do not show spontaneous activity, expression of IEGs does not change in CA1.

These findings demonstrate for the first time an increased potentiality of the VH network for molecular reorganization associated with the spontaneous development of a physiological pattern and may provide a clue to mechanisms that underlie the regulated emergence of SPWs along the longitudinal axis of the hippocampus.

We find that during the first 20-30 minutes of tissue maintenance in vitro, detectable spontaneous SPWs can be recorded in VH slices. Although statistically significant changes in mRNA expression were detected from one hour of tissue maintenance, increases in expression appear to start already during the first fifteen minutes. Therefore, it is difficult to draw a safe conclusion on the specific time relationship between the two phenomena. The fact that increases in mRNA expression were also observed in DH slices in which there is no spontaneous activity could indicate that the molecular reorganization processes occur independently of the spontaneous activity. However, one of the most striking findings in the present study is that an increase in mRNA expression in the CA1 field of the VH is observed for exactly those subunits (i.e. α1/β2/γ2) constituting the most abundant GABA_A_ receptor subtype in the brain including hippocampus (Nusser et al. 1996; Benke et al. 2004). Importantly, the increase in mRNA expression of these subunits is followed by a clear increase in GABA_A_ receptors in the VH suggesting that the molecular reorganization in VH is selective and likely triggered by the ongoing occurrence of spontaneous network activity.

### Implications of increased GABA_A_ receptor expression in SPWs

GABA_A_ receptor is a key component for the generation of SPWs (Buzsaki 2015) and is subject to neuronal activity-induced plastic changes that could lead to profound functional effects (Vithlani et al. 2011; Lorenz-Guertin and Jacob 2017). For instance, changes in GABA_A_ receptor could make a crucial contribution to the regulation of excitation/inhibition balance (Fritschy 2008) and an accurate balance between excitation and inhibition is required for the generation of SPWs (Giannopoulos and Papatheodoropoulos 2013). In particular, the presence of α1 subunit confers the GABA_A_ receptor with increased inhibitory current upon activation (Lavoie et al. 1997; Vicini et al. 2001; Mortensen et al. 2012). This may have a dual implication for SPWs. First, the time-dependent increase in the expression of a α1-containing receptor is compatible with the gradually augmenting amplitude of SPWs in VH slices since SPWs corresponds to GABA_A_ receptor-mediated hyperpolarizations in CA1 pyramidal cells (Papatheodoropoulos 2008; Wu et al. 2002). Second, increased SPW-associated α1-GABA_A_ receptor-mediated hyperpolarization could significantly assist the functional role of SPWs. Specifically, during a given SPW event only a subset of CA1 pyramidal neurons fire (Papatheodoropoulos 2008; Bahner et al. 2011; Chrobak and Buzsaki 1996), thereby constituting the hippocampal output to neocortical regions and subserving by this way the actualization of information transfer between the two brain regions and the ensuing consolidation of memories in neocortex (Buzsaki 1996, 2015). Because SPW-associated hippocampal output is thought to carry specific information, the firing of CA1 pyramidal cells during SPWs must be selective and accurate (Wilson and McNaughton 1994). Although the precise mechanisms that underlie CA1 pyramidal cell firing during SPWs are obscure, however, recent evidence suggests that inhibition may play a crucial role. Specifically, it has been shown that the firing of CA1 neurons, that represent the hippocampal output during SPWs, occurs while most neurons are hyperpolarized (Papatheodoropoulos 2008; Bahner et al. 2011). Thus, it has been proposed that this general GABAergic hyperpolarization of CA1 pyramidal cells during SPWs represent a background mechanism that significantly contributes to the process of selection of “output” CA1 neurons by suppressing the firing of the remaining neurons (Papatheodoropoulos 2008). Consequently, the more effective the inhibition of most neurons is, the more efficient the process of selected cell firing is. Accordingly, assuming that the establishment of SPWs in the hippocampus and the associated transfer of information to neocortex are gradual processes based on the recurrent nature of SPWs (Buzsaki 1989, 2015), it could be speculated that the increased expression of specific GABA_A_ receptor subtype during the gradual development of SPWs in VH slices are functionally interrelated events and that the “teaching” repetitive occurrence of SPWs results in a higher cell hyperpolarization that contributes to efficiency of cell selection process. Also, because α1-containing GABA_A_ receptors on pyramidal cells are located at synapses formed by parvalbumin-expressing basket cells (Klausberger et al. 2002), the increase in α1-GABA_A_ receptors in the CA1 field of VH slices may express a postsynaptic aspect of the mechanisms that underlie the selectively increased activity of parvalbumin-expressing cells during SPWs (Klausberger et al. 2003).

### Network homeostasis and methodological implications

The quantitative changes in the GABA_A_ receptor in SPW-generating VH slices may also represent a homeostatic mechanism that keeps VH network excitability into a physiological range. It is thought that the VH network has an increased intrinsic excitability which under certain conditions can develop into a hyperexcitability state; see references in (Papatheodoropoulos 2018). The generation of SPWs requires a certain increase in excitation (Csicsvari et al. 1999; Giannopoulos and Papatheodoropoulos 2013; Mizunuma et al. 2014) and SPWs have some common features with epileptiform activity including increased synchrony (see (Buzsaki 2015). Their emergence, therefore, poses the risk of the VH network switching to a state of hyperexcitability. In order for the VH network to continue functioning properly, has to balance its endogenous and SPW-associated excitability. Accordingly, the increased number of α1-containing GABA_A_ receptors that accompany the development of SPWs could contribute to the homeostatic regulation of excitability during the emergence and amplification of SPWs. For instance, experimentally induced increase in α1 subunit expression slows down the development of epileptic seizures in hippocampus (Raol et al. 2006). Interestingly, we have noted that the incidence of epileptiform discharges induced in magnesium-free medium is dramatically reduced in VH slices but not in DH slices after 6-8 hours of tissue maintenance (Moschovos and Papatheodoropoulos, unpublished observations). Thus, the present results might be especially relevant for experimental studies of epilepsy given the crucial role that GABA_A_ receptor plays in regulating excitation/inhibition balance and therefore in determining the occurrence of epileptiform activity (Fritschy 2008; Scharfman 2007). Further, studying the mechanisms of GABA_A_ receptor regulation may provide important insights into the molecular underpinnings of physiological conditions that regulate excitability and therefore could help developing treatments that can alleviate pathological conditions like epilepsy, perhaps avoiding the burden of side effects associated with continuous classical pharmacological treatments (Palma et al. 2017). It could be also noted here that the time course of changes in GABA_A_ receptor subunits found in the present study is similar to the rapid adaptive changes that occur in the hippocampal circuitry in response to intense synchronized network activity (Casanova et al. 2013).

### Implications of IEGs expression for SPWs function

A fundamental expression of molecular plasticity is the ability for flexibly regulating gene expression in order neurons and neuronal circuits are effectively adapted to current functional needs (Tsukamoto-Yasui et al. 2007; Miyashita et al. 2008; Minatohara et al. 2016; Buckby et al. 2006; Ziv and Brenner 2018; Turrigiano 2012). In the present study we found that full development of SPWs in the CA1 field of VH slices is accompanied by increased expression of IEGs in this hippocampal field. Remarkably, the expression of IEGs did not change in the CA1 field of DH slices though dentate gyrus and CA3 field of the same slices showed increased IEGs expression.

IEGs are categorized into regulatory transcription factors, like c-fos and zif-268 that control the expression of other genes and effectors, like Arc, that directly influence cellular processes (Kubik et al. 2007; Minatohara et al. 2016) It is widely believed that IEG expression plays a critical role in transforming activity in neural circuits into lasting changes in synaptic activity underlying long-term memory (Minatohara et al. 2016; Diekelmann and Born 2010) Thereby, IEG represent particularly reliable markers of neuronal activity and the induction of IEGs occurs within minutes of increased activity in neuronal networks (Diekelmann and Born 2010). Furthermore, IEGs expression that results in synaptic modifications could contribute to the appearance and the properties of spontaneous activity (Mizunuma et al. 2014). Thus, spontaneous network activity and molecular/synaptic modifications are reciprocally connected (Bains et al. 1999). Importantly, IEG expression is rapidly and selectively upregulated by behavioral experience and neuronal activity in subsets of neurons in specific brain regions which are associated with learning and memory (Kubik et al. 2007; Kim et al. 2018; Minatohara et al. 2016). It is particularly interesting that spontaneous activity in neuronal networks can induce reorganization of synaptic connections, thereby continuously updating the network status (Tsukamoto-Yasui et al. 2007), as it virtually happens in the CA1 circuitry of VH slices, where the increase in amplitude of SPWs is associated with specific molecular and synaptic reorganization. Besides, a basic function ascribed to SPWs is to strengthen synaptic connections in the cell assembly which generates this activity, by virtue of their repetitive occurrence in stages following behavioral experience, thereby leading to establishment of memory trace inside hippocampal circuitry (Buzsaki 1989, 2015).

Several studies have previously reported regional and temporally selective IEG activation patterns (Kubik et al. 2007). Recently aggregated evidence has made it clear that there is a marked differentiation in both constitutive and behavior-induced transcriptional and translational activity along the axis (Sotiriou et al. 2005; Pandis et al. 2006; Gusev et al. 2005; Leonardo et al. 2006; Thompson et al. 2008; Dong et al. 2009; Snyder et al. 2011; Cembrowski et al. 2016; Lee et al. 2017; Chawla et al. 2018; Floriou-Servou et al. 2018). The present results extend this evidence indicating that physiological activity patterns can significantly shape transcriptional heterogeneity, which therefore may further support functional heterogeneity along the hippocampus long axis.

## Conclusions

Summarizing, the present results show that the development of the physiological network activity of SPWs under normal conditions in the VH but not in the DH is associated with a selective increase in the expression of a specific GABA_A_ receptor subtype (α1/β2/γ2) and a selective increased expression of IEGs specifically in the VH field where SPWs are generated. The different gene-expression profiles in DH and VH may result from distinct functional requirements and endow the two hippocampal segments with distinct functional properties. The molecular changes may endow local VH network with specific properties that assist the strengthening of SPWs as was indeed observed. We conclude that the generation of SPWs and the specific molecular reorganization observed in the VH are potentially linked between each other and may represent physiological processes strictly related to the functional roles of SPWs. This evidence suggests that dynamic tuning of the molecular organization may importantly contribute to the functional segregation/heterogeneity seen along the hippocampus. Also, the *in vitro* maintenance of acute hippocampal preparations can provide a methodological powerful tool for investigating the mechanisms underlying the dynamic relationships between neuronal activity and molecular reorganization along the hippocampus aiming at revealing important aspects of the mechanisms underlying the distinct emergent functional properties along the longitudinal axis of the hippocampus.

## Acknowledgements

The study was supported by resources of the Medical School of the University of Patras.

